# The role of B cells in immune cell activation in polycystic ovary syndrome

**DOI:** 10.1101/2023.01.26.525671

**Authors:** Angelo Ascani, Sara Torstensson, Sanjiv Risal, Haojiang Lu, Gustaw Eriksson, Congru Li, Sabrina Teshl, Joana Menezes, Katalin Sandor, Claes Ohlsson, Camilla I Svensson, Mikael C.I. Karlsson, Martin Helmut Stradner, Barbara Obermayer-Pietsch, Elisabet Stener-Victorin

**Affiliations:** Department of Internal Medicine Medical University of Graz, Austria; Department of Physiology and Pharmacology, Karolinska Institutet, Stockholm, Sweden; Centre for Bone and Arthritis Research, Department of Internal Medicine and Clinical Nutrition, Institute of Medicine, Sahlgrenska Academy, University of Gothenburg, Gothenburg, Sweden; Department of Drug Treatment, Sahlgrenska University Hospital, Region Västra Götaland, Gothenburg, Sweden; Department of Microbiology, Tumor and Cell Biology, Karolinska Institutet, Stockholm, Sweden

**Keywords:** polycystic ovary syndrome, B cells, immunology, inflammation

## Abstract

Variations in B cell numbers are associated with polycystic ovary syndrome (PCOS) through unknown mechanisms. Here we demonstrate that B cells are not central mediators of PCOS pathology and that their frequencies are altered as a direct effect of androgen receptor activation. Hyperandrogenic women with PCOS have increased frequencies of age-associated double-negative B memory cells and increased levels of circulating immunoglobulin M (IgM). However, the transfer of serum IgG from women into wild-type female mice induces only an increase in body weight. Furthermore, RAG1 knock-out mice, which lack mature T- and B cells, fail to develop any PCOS-like phenotype. In wild-type mice, co-treatment with flutamide, an androgen receptor antagonist, prevents not only the development of a PCOS-like phenotype but also alterations of B cell frequencies induced by dihydrotestosterone (DHT). Finally, B cell-deficient mice, when exposed to DHT, are not protected from developing a PCOS-like phenotype. These results urge further studies on B cell functions and their effects on autoimmune comorbidities highly prevalent among women with PCOS.

**Summary:** Androgen receptor activation alters B cell frequencies and functionality as the transfer of human PCOS IgG increase weight in female mice. Lack of B cells does not protect from the development of a PCOS phenotype, suggesting an unrecognized role for B cells in PCOS autoimmune comorbidities.

**Graphical Abstract:** 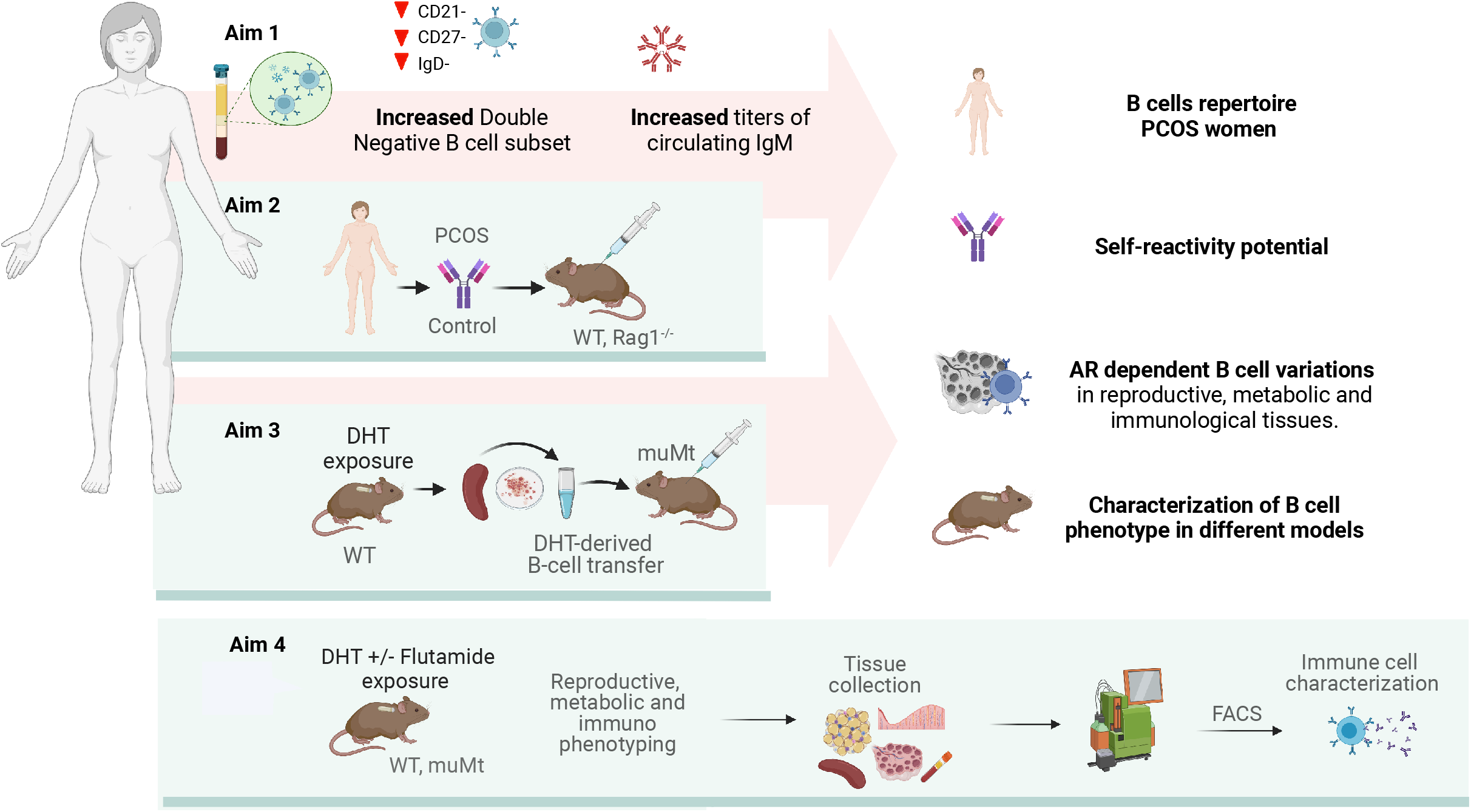

## Introduction

Polycystic ovary syndrome (PCOS) is the single most common endocrine-metabolic disorder affecting 5-18% of women in reproductive age worldwide^1^. As a multifactorial disorder with no clearly defined etiology, PCOS is diagnosed based on the criteria of hyperandrogenism, oligo-anovulation and polycystic ovarian morphology (PCOM)^1^. The syndrome is characterized by chronic low-grade inflammation and potential autoimmune sequelae^2^, further aggravated by obesity. Indeed, women affected with PCOS are at increased risk for type 2 diabetes mellitus (T2D)^3^ with a number of studies indicating also a higher prevalence of autoimmune thyroid disease (AITD), and particularly hypothyroidism^2^. In support of this observation, serologic parameters of autoimmunity, such as anti-histone and anti–double-stranded DNA antibodies, are relatively high among these women^4^. However, attempts to identify an autoimmune cause of PCOS have been uninformative^5–7^. Hyperandrogenemia is a hallmark feature of PCOS that plays a key role in the pathophysiology and seems to be directly related to disease severity^1,8^. Although chronic inflammation and altered immune function have been proposed to play a role in the pathogenesis of PCOS and T2D, it remains unknown whether the observed immune responses and autoimmune alterations in women with PCOS are a cause or consequence of hyperandrogenemia^9^.

The notion of B cells exacerbating metabolic disease has been known for over a decade, both in the pathogenesis of diabetes^10^ as well as in obesity-associated insulin resistance^11^. Although being an extreme model, biological variations in mu heavy chain knockout mice (MuMt^-^; B^null^), which fail to produce mature B cells, have contributed to this knowledge^11^. Notably, muMt^-^ mice reconstituted with B cells derived from mice with diet-induced obesity (DIO) develop an impaired glucose tolerance. However, when transferring immunoglobulin G (IgG) from diet-induced obesity mice to muMt mice, systemic inflammatory changes were noticeable only when the recipient mice were exposed to an high-fat diet, suggesting that metabolic effects stemming from B cells may require exposure to a prior stimulus as a determinant of reaction, via induced conditioning or induction of target autoantigens^11^.

These observations led us to investigate whether the hyperandrogenic hormonal milieu in PCOS could have a predominant role in B cell fate, as the androgen receptor (AR) is expressed both in immune organs as well as on precursors and some mature immune cells, potentially implicating various levels of susceptibility^12^. Testosterone is also an indirect regulator of the cytokine B cell activating factor (BAFF), also known as TNFSF13B, an essential survival factor for B cells^13^ which has been shown to be increased in women affected with PCOS^14^. This may be a plausible candidate mechanism as excessive BAFF levels allow for the survival of autoreactive cells and autoantibody production^15^. Recent research has shown that the proportions and activity of peripheral B cells in women with PCOS are increased^14^, though it remains unclear whether B cells alone are the main inflammatory drivers of PCOS pathogenesis and if hyperandrogenemia through AR activation may lead to the acquisition of their unique characteristics. When examining potential autoreactive B cells in PCOS it is important and clinically meaningful to discern the role of androgen exposure alone on the immune system from that of obesity-deriving inflammation. Plasma of individuals with obesity has been shown to be enriched in IgG antibodies with anti-self-reactivity, which have been positively associated with blood frequencies of double negative (DN) B cells, considered the most pro-inflammatory B cell subset^16^. How androgen exposure alone, through androgen receptor activation, affect B cells and their function in a PCOS-like mice model exhibiting a reproductive and metabolic phenotype^17^, remains unknown.

In this study, we investigate the role of B cells in the underlying inflammation of PCOS by assessing the effect of hyperandrogenism on B cell populations and whether B cells are contributing to the pathology. For this purpose, in line with previous data coupling increased B cells numbers (CD19^+^) with PCOS, we first aimed to define which main B cell lineages are affected in hyperandrogenic women with PCOS. Next, we assessed if B cells with self-reactive potential may have a causal effect on both the development of a PCOS-like phenotype, including metabolic dysfunction, and the immune profile in mice by transferring IgG from women with PCOS. To further study if B cell frequencies are altered in reproductive, metabolic, and immunological tissues, major variations of B cell subsets were analyzed in a dihydrotestosterone (DHT)-induced PCOS-like mouse model. Furthermore, we investigated whether these DHT-induced alterations are a result of androgen receptor activation by simultaneous administration of flutamide, an androgen receptor antagonist. Finally, we questioned whether B cell deficient muMt^-^ mice are protected from developing PCOS traits when exposed to DHT.

Here, we demonstrate that AR activation is a direct modulator of B cell frequencies in PCOS pathogenesis. We show that the transfer of circulating IgG from women with PCOS disrupts B cell proportions and causes an increased body weight and sex steroid imbalance in female WT mice. Collectively, our data support a model wherein activation of B cells promotes the development of a PCOS-like metabolic phenotype in mice. We also suggest caution towards therapeutically targeting CD19^+^ cells as DHT-exposed B cell-deficient mice develop a PCOS-like phenotype, showing that a lack of B cells is not protective and reiterate the need for broader studies on alterations of the immune system within the complex hormonal frame of PCOS, including activation of T cells and tissue-resident immune cells.

## Results

### Androgens are associated with altered B cell frequencies and immunoglobulin M increase in women with PCOS

As alterations of B cell frequencies have previously been shown in women with PCOS^14^, we first characterized main B cell lineages and subpopulations based on pan B cell surface marker CD19 in the serum of 15 hyperandrogenic women with PCOS and of 22 women without PCOS (controls). Women with PCOS fulfilled all three Rotterdam Criteria, displaying oligo-/amenorrhea, hirsutism, and polycystic ovarian morphology. Women with PCOS were younger than controls, with median ages of 26 and 36, respectively, with no difference in BMI, but with significantly higher total testosterone and androstenedione with elevated total triglycerides, and reduced HDL-cholesterol (**Table S1**). CD19^+^ B cell memory populations were phenotypically analyzed based on surface markers IgD and CD27. Our initial assessment showed a remodeling of B cell repertoire in women with PCOS compared to controls. The frequency of age-associated DN B memory cells lacking surface expression of CD27 and IgD was significantly higher in women with PCOS (**Fig. 1a**), with declined “innate-like” unswitched CD27^+^IgD^+^ B memory cells (**Fig. 1b**). While naïve B cells populations did not differ among study groups (**Fig. 1c**), switched CD27^+^ IgD^-^ were increased in women with PCOS (**Fig. 1d**), which may directly affect unswitched B cells frequency variance. We did not find direct evidence of activation of DN B cells among women affected by PCOS. Analysis of surface marker CD38, generally expressed on antibody-secreting plasma cells, proved similar values in both groups (**Fig. 1e**). Expression of CD86, a co-stimulatory molecule which usually is upregulated following activation of B cells and in turn can activate T cells, did not differ significantly either (**Fig. 1f**). When assessing circulating serum antibodies, immunoglobulins M (IgM) were higher in hyperandrogenic women with PCOS (**Fig. 1g**) exhibiting high testosterone and increased Free Androgen Index (**Fig. 1h-i**) compared to controls with similar BMI (**Fig. 1j**). Interestingly, no differences were noted for circulating IgG (**Fig. S1a**) while IgA titers were lower in women with PCOS (**Fig. S1b**).

**Fig. 1.**
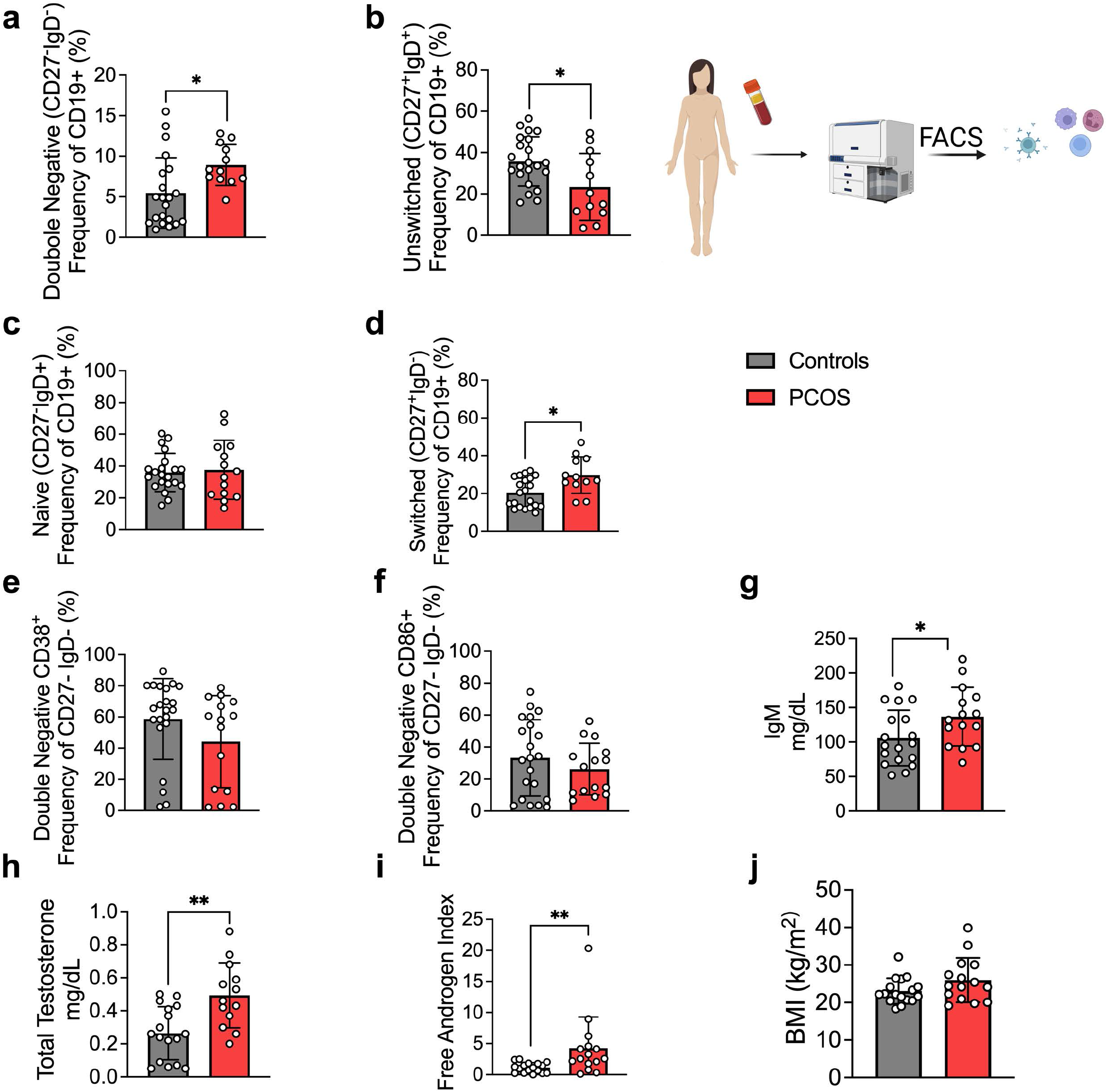
B cell frequencies and immunoglobulin M variations in women with Polycystic Ovary Syndrome. **a.** Total CD19+ Double Negative B cells (CD27-IgD-) **b.** Total Unswitched B cells (CD27+ lgD+) **c.** Total Naive B cells (CD27-IgD+) **d.** Total Switched B cells (CD27+IgD-) **a-d** Total CD19+ populations controls n=22; PCOS n=15; **e-f** Expression on Double Negative B cells respectively of the surface markers CD38 and CD86 **g.** Circulating IgM titers (controls n=18; PCOS n=15) **h.** Total testosterone **i.** Free Androgen Index (FAI) **j.** Body Mass Index. All bars indicate means, error bars SD, circles represent human individuals. Unpaired Student’s t-test for analysis of naive, unswitched and DN CD86+ B cells, total testosterone, and BMI. Mann–Whitney test for all other B cell frequencies, antibody titers and FAI. *P<0.05, **P<0.01, ***P<0.001.

These data support the hypothesis that women with PCOS and hyperandrogenemia have an altered B-cell frequency linked to alterations in IgM antibody production. However, higher disease activity was not explained by increased DN B lymphopoiesis.

### Transfer of human derived IgG antibodies results in increased body weight in WT female mice

Clusters of pro-inflammatory age-associated double negative (DN) B memory cells lacking surface expression of CD27, and immunoglobulin D (IgD) have been associated with plasma cell differentiation fate, and while not increasing significantly in numbers, produce higher amounts of IgG on a per cell basis relative to switched memory B cells^18^. Hence, to assess if PCOS may have an underlying autoimmunological effector component, we investigated a possible role for IgG in PCOS systemic inflammation. IgG antibody extracted from serum of women with PCOS diagnosed as phenotype A fulfilling all three Rotterdam Criteria, displaying oligo-/amenorrhea, hirsutism, and polycystic ovarian morphology and of healthy controls (**Table S2**) were purified and pooled, then transferred intraperitoneally (i.p.) into wild-type (WT) mice. Following the same procedure, IgG deriving from hormonally healthy women was equally purified, then pooled and transferred into similar age and weight-matched WT mice. There were no differences in ovulatory cycles between controls (**Fig. 2a**) and mice receiving PCOS IgG (**Fig. 2b**). Three weeks post IgG transfer, mice receiving IgG from women with PCOS increased in body weight compared to controls (**Fig. 2 c**). Body composition assessment showed no difference in proportion of fat or lean mass between the groups (**Fig. 2d**). Interestingly, as an effect of human PCOS IgG transfer, recipient mice had also altered subsets of B lymphocytes in blood, ovary, and visceral adipose tissue (VAT). Circulating DN B memory cells were increased (**Fig. 2e**) while blood naïve cells were reduced (**Fig. 2f**), resembling the B cell distribution described in donor women with PCOS. Among the DN B cells, DN1 CD21^+^ subset was the main circulating subpopulation in the blood of mice that received IgG from women with PCOS (**Fig. 2g**). VAT tissue had higher frequencies of effector IgM^+^IgD^+^CD27^+^ “double positive” unswitched B cells (**Fig. 2h**) while activated switched IgM^+^IgD-CD27^+^ were increased in ovarian tissue (**Fig. 2i**). Analyzing circulating sex steroids in these mice, estrogens were altered with an increase in estrone (**Fig. 2 j**) and a trend of higher estradiol (**Fig. 2 k**) with no difference in androgens or progesterone (**Fig. 2 l-n**).

**Fig. 2.**
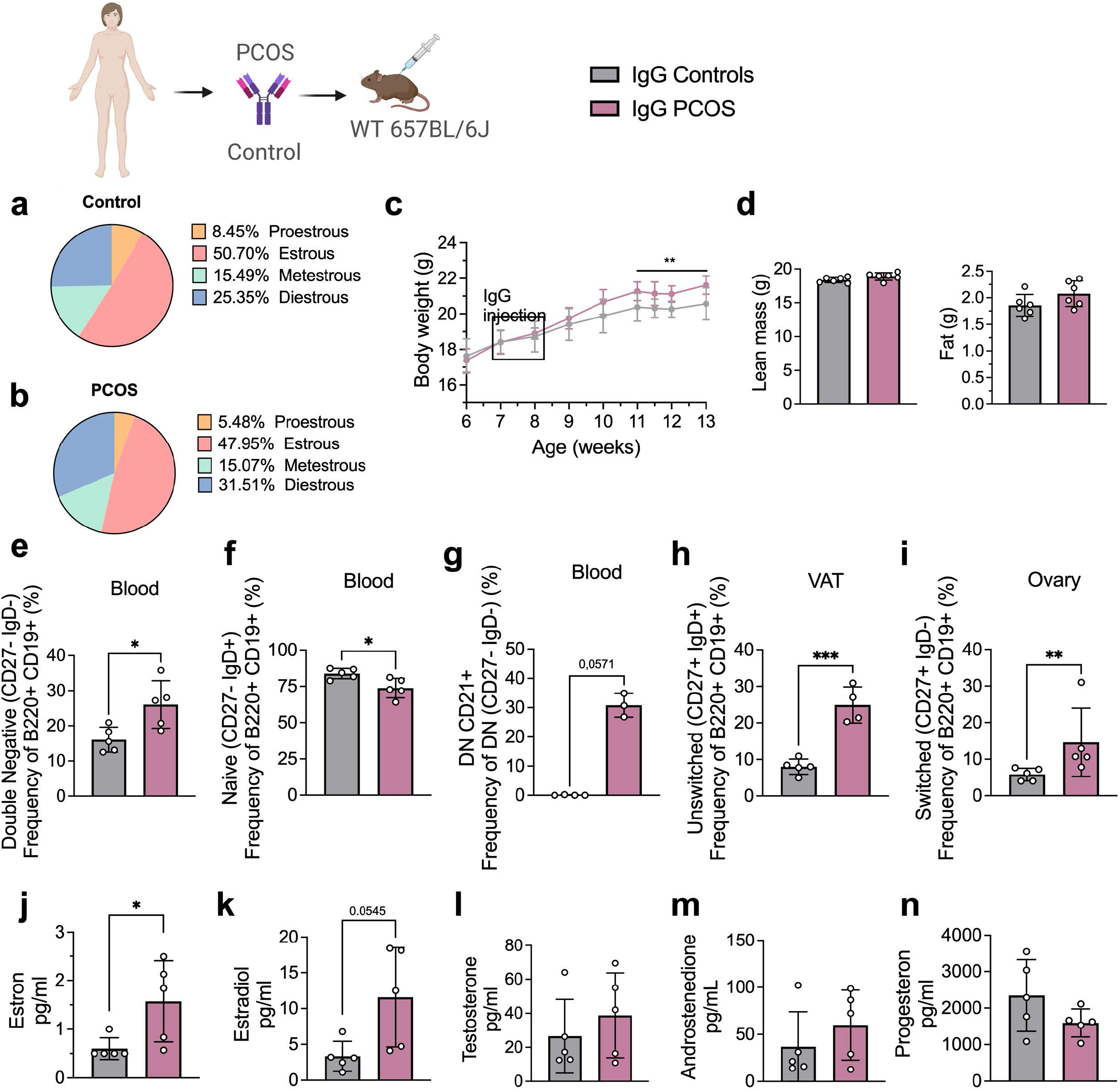
IgG Transfer to WT mice. **a.** Estrous cycles in mice receiving Control IgG **b.** Estrous cycles in mice receiving PCOS IgG **c.** Weekly Body Weight recordings **d.** EcoMRI results for body fat and lean mass composition **e.** Double Negative B cells in blood **f.** Naive B cells in blood **g.** DN CD21+ B cells in blood **h.** Unswitched B cells in VAT **i.** Switched B cells in Ovary **j.** Testosterone **k.** Androstenedione **l.** Estradiol **m.** Estron **n.** Progesterone. All bars indicate means, circles represent individual mice. Unpaired Student’s t-test for analysis of EchoMRI results and all B cell frequencies except DN CD21+ and switched; two-way ANOVA with Sidak’s post-hoc test for analysis of weekly BW recordings; Mann-Whitney test for analysis of DN CD21+, switched, estron, testosterone and androstenedione, Welch’s t-test for analysis of estradiol and progesteron. *P<0.05, **P<0.01, ***P<0.001.

As B cell functions are influenced by other lymphocyte populations, especially T cells, and vice versa, we aimed to assess if T cells are modulating this inflammatory effect deriving from IgG-induced disease. Human IgG deriving from PCOS and control cohorts, was first purified, then pooled into separate groups, and transferred into 10-week-old Rag1 KO^-/-^ mice, which lack of mature T- and B-cells. Three weeks post i.p. IgG transfer, RAG1 KO^-/-^ mice failed to develop any PCOS-like phenotype, contrasting previous transfer into WT mice, suggesting that the pathophysiological mechanism inducing immune and metabolic disruption such as body weight alteration may necessarily involve other lymphocytes to fully promote impairment of metabolic parameters.

### Altered B cell frequencies are replicated in a DHT-induced PCOS-like mouse model and seen in reproductive, metabolic and immunological tissues

To investigate androgen-mediated regulation of B cell phenotypes, in particular DN B memory cells as well as circulating antibody titers, in tissues other than blood, we used the well-established peripubertal DHT-induced PCOS mouse model^19^. Peripubertal female mice were subcutaneously implanted with a silastic pellet containing 4 mm of DHT and develop PCOS-like traits with reproductive and metabolic dysfunction without increase in fat mass^20^. Control mice received an empty, blank implant. To investigate whether any phenotypic differences are driven by AR activation, a third group received, in addition to the DHT implant, a continuously releasing flutamide pellet, an AR antagonist. Two separate cohorts of these three experimental groups were phenotypically characterized at 13 and 16 weeks of age respectively.

After 4 weeks of continuous DHT exposure, mice had developed a reproductive PCOS-like phenotype, exhibiting disrupted estrous cycles, arrested in diestrus or metestrus (**Fig. 3a-b**), and longer anogenital distance (**Fig. 3d**). Co-treatment with flutamide prevented the development of these phenotypes (**Fig. 3c-d**). DHT-exposed mice gained body weight (**Fig. 3e**), not in fat mass (**Fig. 3f**) but rather by increased lean mass (**Fig. 3g**) compared to controls and mice co-treated with flutamide. At 12 weeks of age, no clear sign of impaired glucose homeostasis was noted during oral glucose tolerance test (oGTT) when compared to controls (**Fig. 3h**), although DHT-exposed mice display a trend of higher fasting glucose (**Fig. 3i**).

**Fig. 3.**
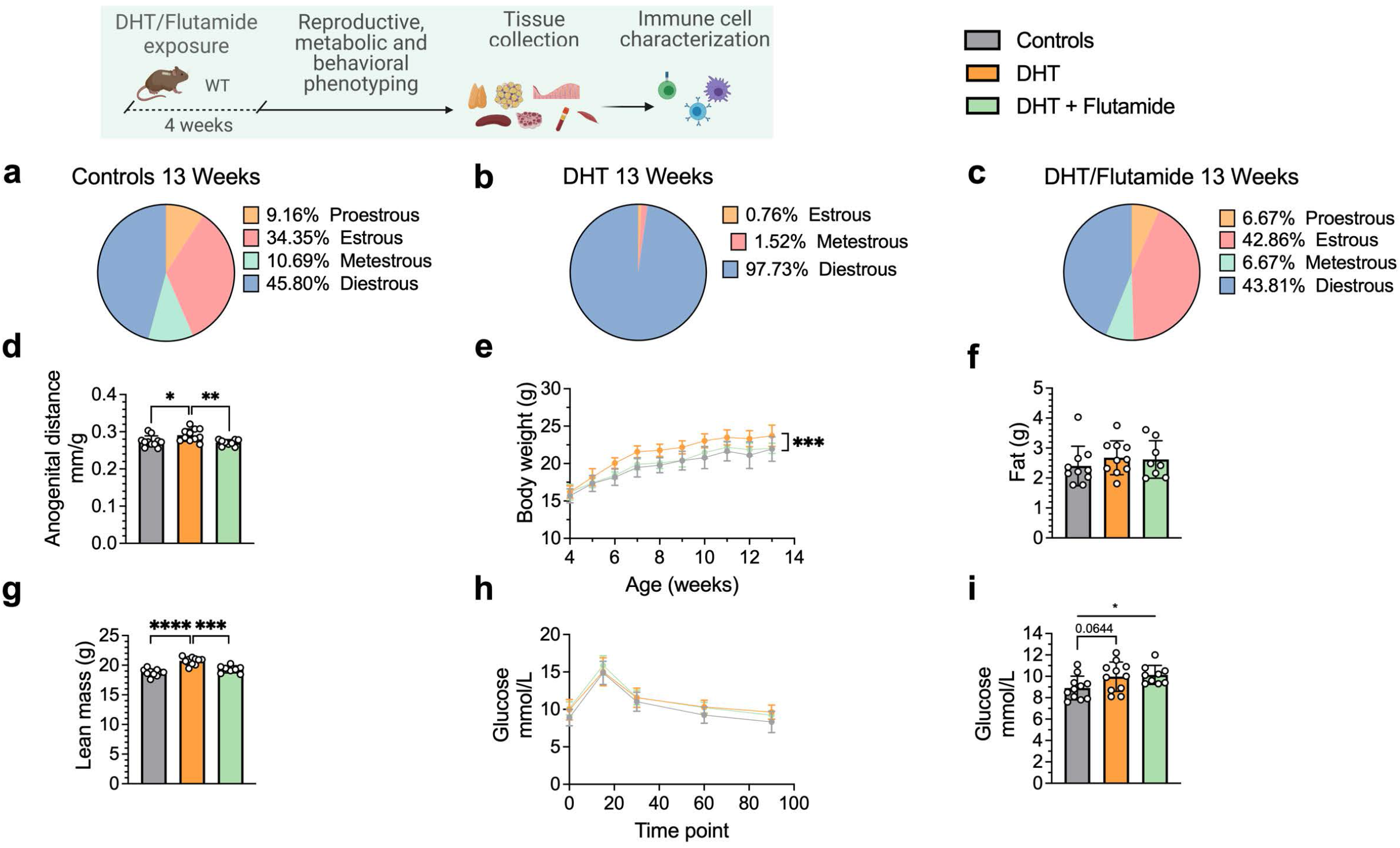
DHT-induced PCOS-like mouse model phenotypic study at 13 weeks of age. **a.** Estrous cycles in WT control mice **b.** Estrous cycles in mice receiving DHT pellet implant **c.** Estrous cycles in mice receiving DHT pellet and Flutamide implant **d.** Normalized anogenital distance **e.** Weekly body weight **f.** EchoMRI record of fat body composition **g.** EchoMRI record of lean body composition **h.** OgTT **i.** Fasting glucose. All bars indicate means, circles represent individual mice. Unpaired Student’s t-test for analysis of anogenital distance difference between groups, as well as EchoMRI results and fasting glucose; two-way ANOVA with Sidak’s post-hoc test for analysis of weekly BW recordings and blood glucose throughout the study. *P<0.05, **P<0.01, ***P<0.001.

At 16 weeks of age, DHT-exposed mice with a PCOS-like phenotype have an equally disrupted estrous cycle when compared to controls (**Fig. 4a**), arrested in diestrus or metestrus (F**ig. 4b**) and longer anogenital distance (**Fig. 4d**). These effects were prevented by co-treatment with flutamide as in previous experiment (**Fig. 4c-d**). DHT-exposed mice weighed more (**Fig. 4e**), an effect due to higher lean mass (fat mass **Fig. 4f**; lean mass **Fig. 4g**), and had impaired glucose homeostasis, displaying an overall impaired glucose uptake during oGTT (**Fig. 4h**) and a higher fasting glucose and when compared to controls (**Fig. 4i**).

**Fig. 4.**
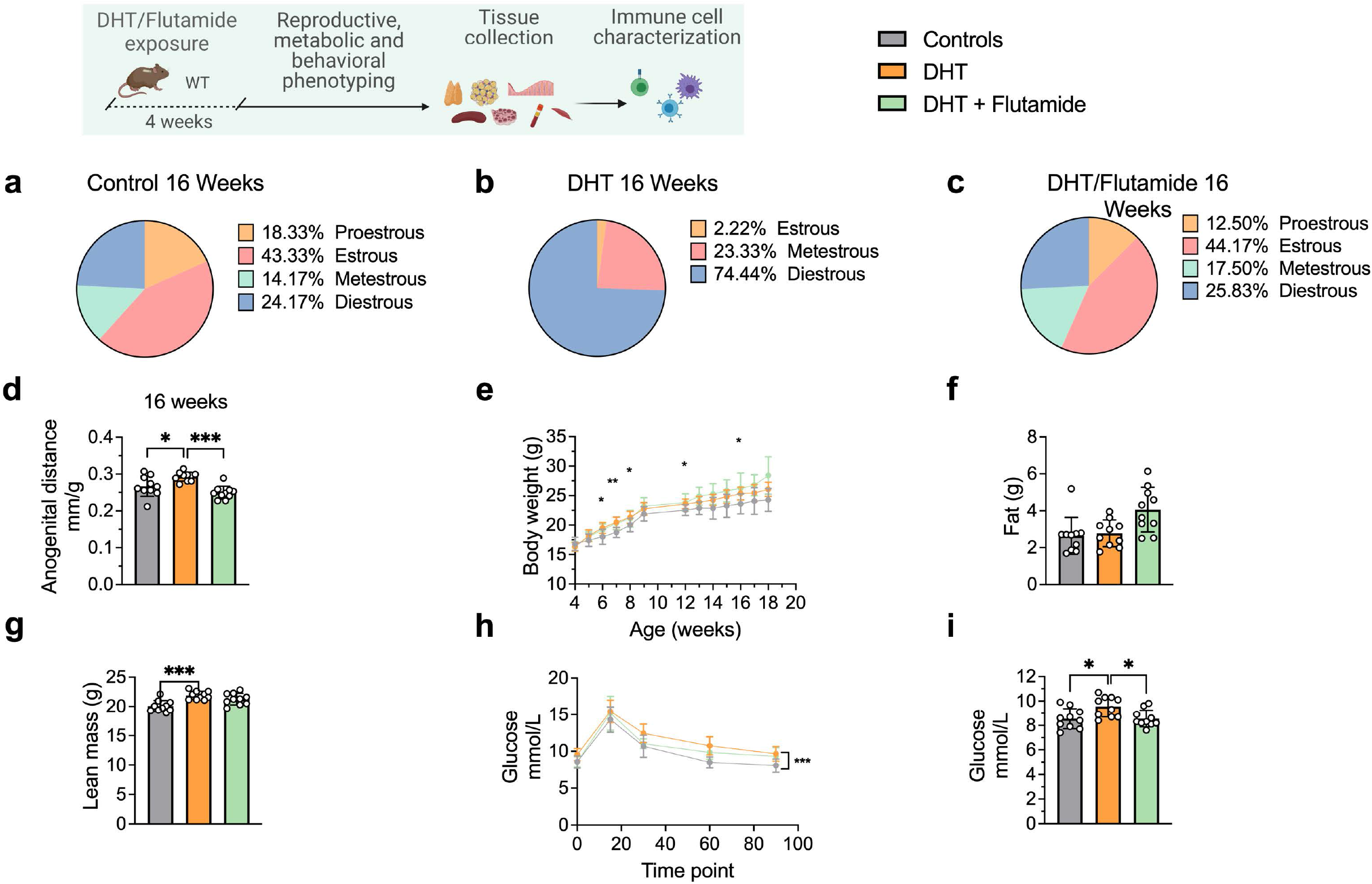
DHT-induced PCOS-like mouse model phenotypic study at 16 weeks of age. **a.** Estrous cycles in WT control mice **b.** Estrous cycles in mice receiving DHT pellet implant **c.** Estrous cycles in mice receiving DHT pellet and Flutamide implant **d.** Anogenital distance normalized to body weight **e.** Weekly body weight **f.** EchoMRI record of fat body composition **g.** EchoMRI record of lean body composition **h.** OgTT **i.** Fasting glucose. All bars indicate means, circles represent individual mice. Unpaired Student’s t-test for analysis of anogenital distance difference between groups, as well as EchoMRI results and fasting glucose; two-way ANOVA with Sidak’s post-hoc test for analysis of weekly BW recordings and blood glucose throughout the study. *P<0.05, **P<0.01, ***P<0.001.

Frequencies of B memory cells in 13-week-old DHT-exposed mice were disrupted compared to controls, particularly in blood. Circulating CD19^+^ DN memory cells were lower compared to controls (**Fig. 5a**). CD19^+^ naïve B cells were increased in the blood of DHT-exposed mice (**Fig 5b**). When analyzing B cell distribution at 20 weeks of age, CD19^+^ DN B cells were increased in the spleen of DHT-exposed mice (**Fig. 5c**) while frequencies of naïve B cells were decreased (**Fig. 5d**), an effect that was reversed when co-treated with flutamide. Overall, ovarian tissue was the most affected tissue. DHT-exposed mice had decreased proportions of both DN and naïve B cells within the ovaries (**Fig 5e-f**), a similar trend as seen in blood-deriving cells of mice at 13 weeks of age. Ovaries of DHT-exposed mice were characterized by an increased frequency of IgM^+^IgD^+^CD27^+^ “double positive” unswitched B cells (**Fig. 5g**). Among the DN cells, DHT-exposed mice displayed a trend, suggesting an increase of DN CD21^+^ populations in the ovaries compared with controls and mice co-treated with flutamide (**Fig. 5h**), an effect previously observed in the blood of mice receiving IgG from women with PCOS. A similar increase was noted among naïve B cells of DHT-exposed mice, with a trend suggesting increased proportions of CD19^+^ naïve B cells expressing CD21^+^ in the ovaries (**Fig. 5i**) and endometrium (**Fig. 5j**), as well as spleen (**Fig. 5k**) and VAT (**Fig. 5l**). These trends were reversed by co-treatment with flutamide in ovary and endometrium.

**Fig. 5.**
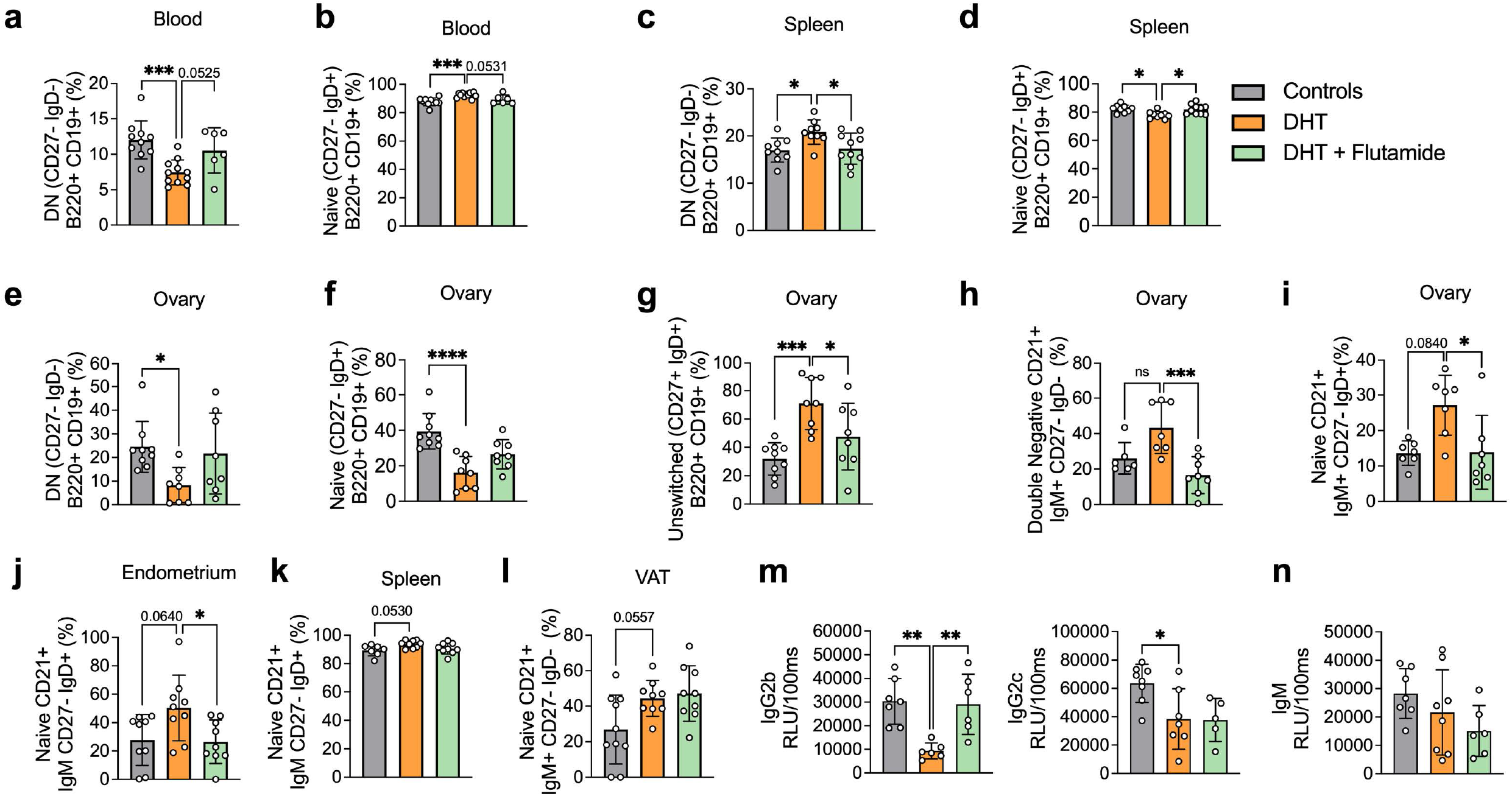
DHT-induced PCOS-like mouse model B cell frequencies. a. Blood DN B cells in 13 week old mice b. Blood Naive B cells in 13-week-old mice c. Spleen DN B cells in 20 week old mice d. Spleen Naive B cells in 20 week old mice e. Ovary DN B cells in 20 week old mice f. Ovary Naive B cells in 20 week old mice g. Ovary Unswitched B cells in 20 week old mice h. Ovary DN CD21+ B cells in 20 week old mice i. Ovary Naive CD21+ B cells in 20 week old mice j. VAT Naive CD21+ B cells in 20 week old mice k. Spleen Naive CD21+ B cells in 16 week old mice l. Endometrium Naive CD21+ B cells in 20 week old mice m. Circulating IgG titers in 20 week old mice. n. Circulating IgM titers in 20 week old mice. All bars indicate means, circles represent individual mice. One-way ANOVA for multiple comparisons of normally distributed data, Kuskal-Wallis test for data that is not normally distributed. *P<0.05, **P<0.01, ***P<0.001.

Collectively, these results point to an inflammatory activity ongoing in the DHT-exposed mice presenting a PCOS-like phenotype, with B cell alterations being a consequence of AR activation as proven by the preventive effect of flutamide cotreatment. There are noticeable differences within the single tissues which require further investigations.

### DHT-induced PCOS-like mice show a distinct IgG profile

In addition to functions deriving from T-cell interaction, B cells regulate immune function via antibody production. Given the altered titers of IgM in women with PCOS, circulating IgM levels as well as IgG isotypes were analyzed in the peripubertal PCOS-like mouse model. DHT-exposed mice, exhibiting elevated levels of circulating testosterone and androstenedione, display reduced levels of IgG2b and IgG2c isotypes (**Fig. 5m**), while no significant differences in IgM levels could be seen (**Fig. 5n**). No differences were found for IgG1 (**Fig. S2a**) nor in IgG3 titers (**Fig. S2b**).

### B cell transfer from DHT-induced PCOS-like mice into B cell deficient mice does not induce a PCOS-like phenotype

To discern the role of B cells in the etiology of PCOS and the development of associated metabolic comorbidities, it was assessed whether transfer of B cells alone from DHT-exposed mice could induce a PCOS-like phenotype in B cell deficient muMt^-^ mice. Splenic B cells from DHT-exposed mice were transferred i.p. in to 6-week-old muMt^-^ B^null^ mice, as they do not produce mature B cells due to the knockout of the mu heavy chain. It is important to note that they have, however, a fully functional T cell compartment. Control muMt^-^ mice received an equal amount of splenic B cells deriving from a control donor group. Two weeks after transfer, the DHT-exposed B cells recipient muMt^-^ mice failed to develop PCOS-like traits. Overall, B cell transfer did not affect the estrous cyclicity (**Fig. 6 a-b**), nor the anogenital distance (**Fig. 6c**). Total body weight did not differ (**Fig. 6d**), nor fat or lean mass (**Fig. 6e**). Fasting glucose was not affected (**Fig. 6f**) nor was glucose tolerance in oGTT testing (**Fig. 6g**). The lack of a PCOS-like phenotype in a B cell reconstituted model with conserved T cell function was not explained by the presence of B cells alone to a hyperandrogenic environment and must therefore be driven by a peripheral mechanism which necessarily also affects function and properties of other immune cells.

**Fig. 6.**
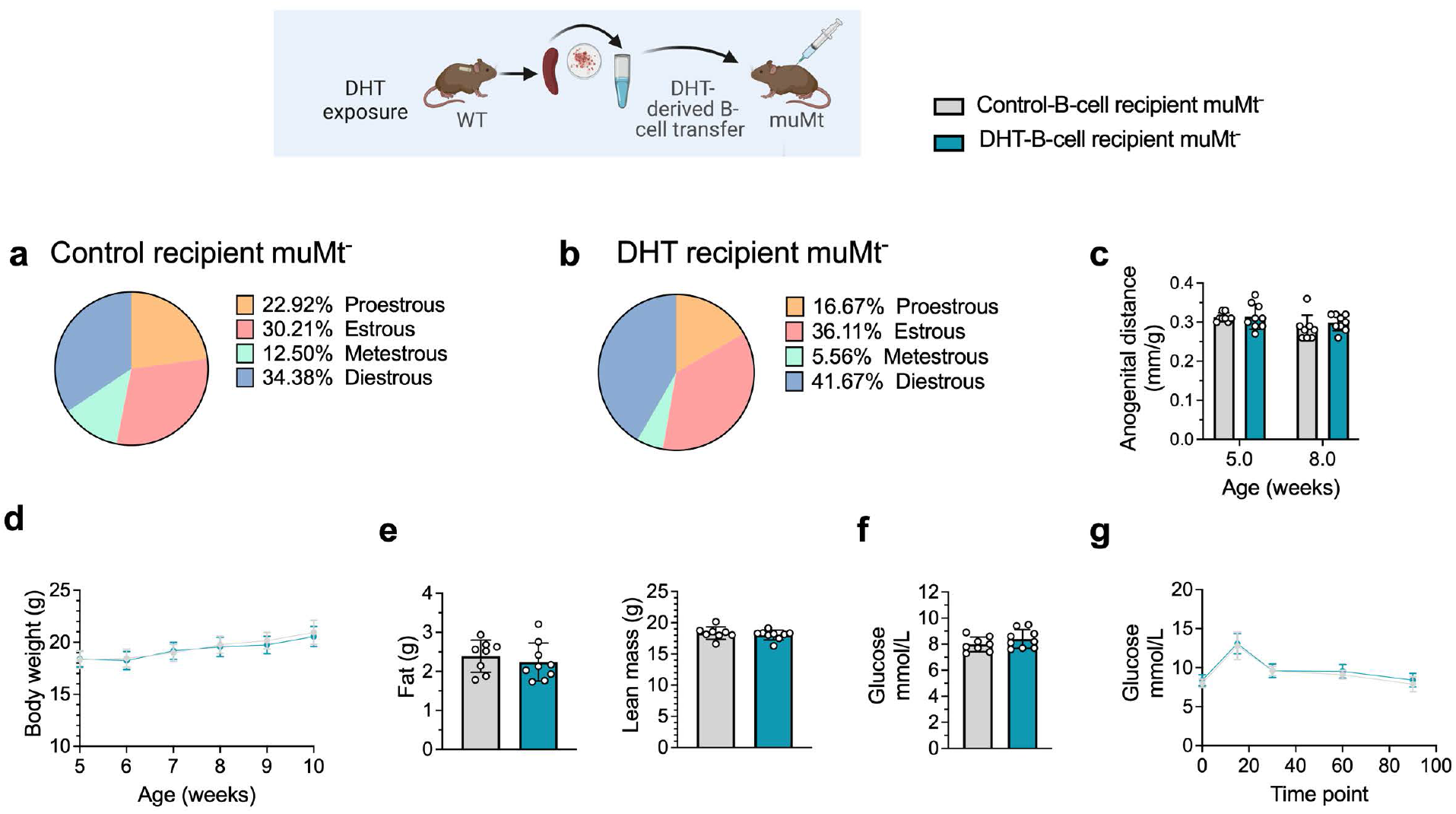
B cell transfer from DHT-induced PCOS-like mice into recipient muMt-B cell-deficient mice. **a.** Estrous cycles in 13-week-old muMt-mice receiving control B cells **b.** Estrous cycles in 13-week-old muMt-mice receiving DHT exposed B cells **c.** Normalized anogenital distance in 13-week-old muMt-recipient mice **d.** Weekly body weight in 13-week-old muMt-recipient mice **e.** EchoMRI record of fat and lean body composition in 13-week-old muMt-recipient mice **f.** Fasting glucose levels. **g.** OgTT in 13-week-old muMt-recipient mice. All bars indicate means, circles represent individual mice. Unpaired Student’s t-test for analysis of anogenital distance difference between groups, as well as EchoMRI results and fasting glucose; two-way ANOVA with Sidak’s post-hoc test for analysis of weekly BW recordings and blood glucose throughout the study; *P<0.05, **P<0.01, ***P<0.001.

### B cell deficiency does not protect from the induction of a PCOS-like phenotype by DHT exposure

To finally assess if B cell deficiency provides a protective effect, 28 days/4-weeks-old muMt^-^ mice, lacking mature B cells, were implanted with a silastic implant containing continuously releasing low-dose DHT. Control mice received a blank pellet. Four weeks after implantation, DHT-exposed B^null^ muMt^-^ developed a clear reproductive PCOS-like phenotype, exhibiting a disrupted estrous cycle (**Fig. 7a-b**), arrested in the diestrus phase, along with longer anogenital distance (**Fig. 7c**). Furthermore, while no difference in body weight was noted amongst the groups at implantation (**Fig. 7d**), DHT-exposed muMt^-^ mice gain higher body weight compared to controls already after one week following implantation (**Fig. 7e**), with increase both in total fat and lean mass (**Fig.7f**). When challenged to oGTT, DHT-exposed muMt^-^ mice exhibited impaired glucose homeostasis, with higher fasting glucose levels (**Fig. 7g**) and higher blood glucose score 90 minutes after administration compared to control (**Fig. 7h**).

**Fig. 7.**
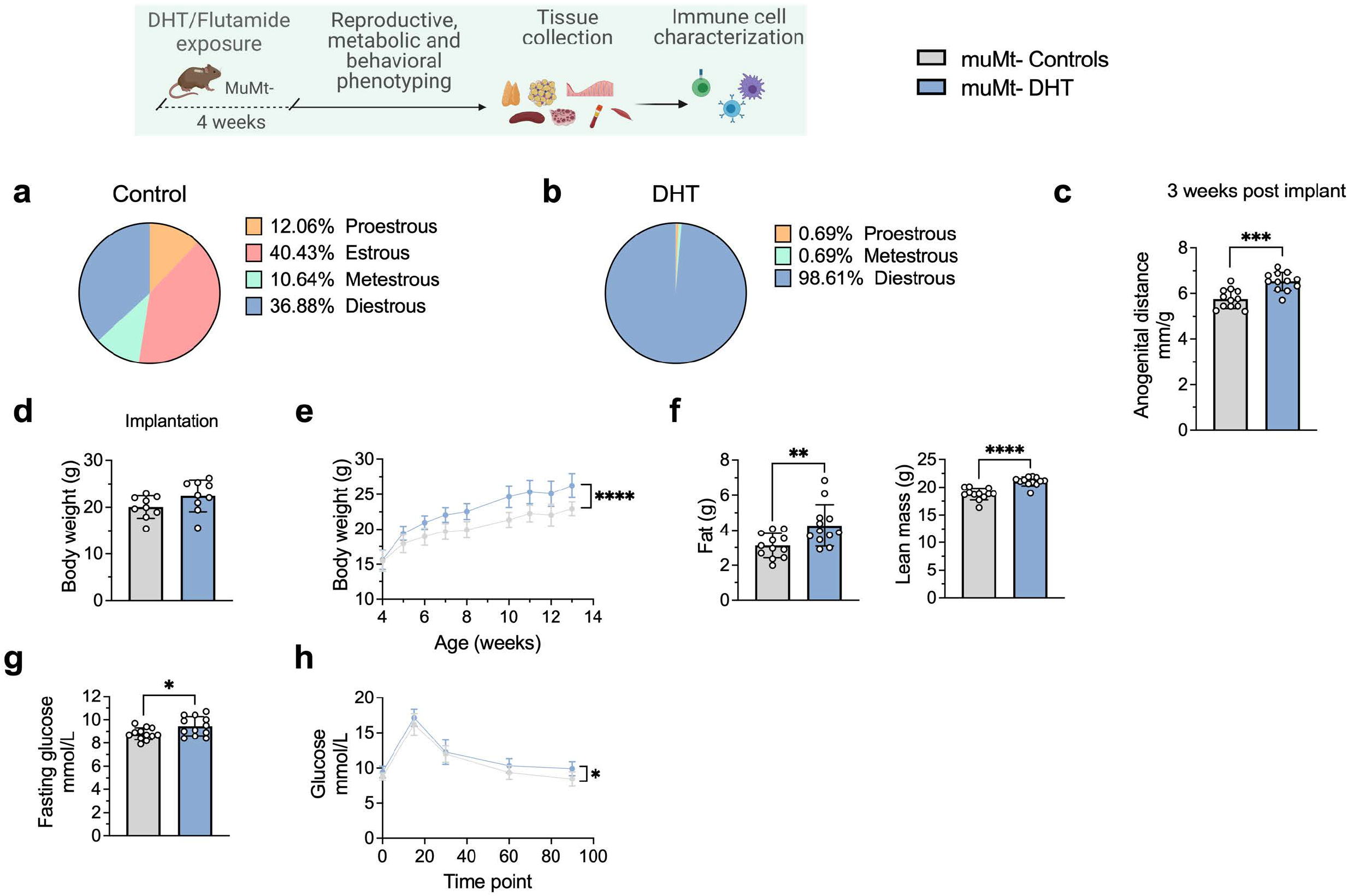
MuMt-DHT-induced mouse model phenotypic study. **a.** Estrous cycles in muMt-control mice **b.** Estrous cycles in muMt-mice receiving DHT pellet implant **c.** Normalized anogenital distance 3 weeks post pellet implantation **d.** Body weight at pellet implantation **e.** Weekly body weight recordings **f.** EchoMRI record of fat and lean body composition **g.** Fasting glucose levels **h.** Oral Glucose Test results. All bars indicate means, circles represent individual mice. Unpaired Student’s t-test for analysis of anogenital distance difference between groups, as well as BW at implantation, fat mass and fasting glucose; Mann-Whitney test for analysis of lean mass; two-way ANOVA with Sidak’s post-hoc test for analysis of weekly BW recordings and blood glucose throughout the study *P<0.05, **P<0.01, ***P<0.001.

These data firstly suggest that although affected in their function, B cells may not be central mediators of glucose metabolism impairment and reproductive dysfunction in PCOS. Furthermore, these mice develop a worsened outcome with increased adipose tissue which was unaltered in previous models, a risk factor of increased disease severity which is aggravated by obesity.

## Discussion

Here we show a distinct link between hyperandrogenemia and abnormal B cell frequencies with circulating antibody titers in women with PCOS, potentially exerting effects systemically via pathogenic IgG antibodies inducing metabolic alterations leading to increased body weight as seen in mice following IgG transfer from these women. We further demonstrate how these highly regulated mechanisms are intrinsically AR activation-dependent. Most importantly, we show that B cells are not the central mediators of systemic inflammation or glucose metabolism impairment in PCOS, as a lack of these lymphocytes does not protect from the induction of a PCOS-like phenotype following DHT exposure. Rather the DHT-exposed muMt^-^ mice display an aggravated phenotype with higher fat mass accumulation.

In line with data combining increased BAFF levels and higher B cell frequencies in PCOS^14^, hyperandrogenic women with PCOS had a significant rearrangement of the B cell repertoire, resulting in higher frequencies of DN B memory cells which notably differ from naïve B cells for their lack of IgD. These age-associated or tissue-based atypical memory B cells have been previously described as autoantibody enriched and poised for plasma cell differentiation^18^. DN B memory cells over-accumulate in chronic infections, autoimmune diseases, and immunodeficiency^21,22^. Based on B cells activation status and antibody-secreting potential, our findings did not support an ongoing proinflammatory activity driven by this specific heterogenous cluster as a major underlying mechanism in PCOS. Markers for B cell activation and T cell costimulatory capacity (CD86), as well as antibody secretion potential (CD38), did not differ from the DN memory B cells of our control population. Moreover, DN B memory cells deriving from women with PCOS were largely IgM-positive, resembling IgM-only cells. Previous studies have suggested this subset to be a major origin of switched-memory B cells^23^. In humans, switched memory B cells have the propensity to differentiate into plasma cells upon reactivation^24^. These results may indicate that PCOS represents a state of prolonged trained immunity, susceptible to secondary triggers of inflammation. Consistent with this finding, by combining both markers IgD and CD27 in our study, we were able to distinguish a decrease in unswitched CD27^+^ IgD^+^ memory B cells accompanied by increased proportions of switched CD27^+^ IgD^-^ memory B cells in women with PCOS. Interestingly, a strong positive correlation was found between circulating IgM prevalence and androgens in women with PCOS, which was not affected by BMI. While increased adiposity is a determining factor in chronic inflammation, PCOS clearly triggers a condition-specific antigenic stimulus causing rearrangement of the humoral immune system.

There are data describing increased serum levels of autoantibodies (e.g., anti-histone and anti–double-stranded DNA antibodies) in PCOS^4^. Additionally, a dose-dependent activation effect has been shown *in vitro* using PCOS-derived purified serum IgG antibodies on the gonadotropin-releasing hormone receptor^25^ (GnRH). In the present study, the transfer of purified and concentrated serum IgG antibodies deriving from women with hyperandrogenic PCOS did increase body weight in recipient mice, although there was no biological variation in ovulatory cycles, a universal PCOS-like trait. Interestingly, we also found a concomitant disruption of B cell frequencies and a sex hormone imbalance among estrogens, partially reflecting the status of PCOS donors. Recipient mice of human PCOS IgG had in fact increased circulating DN memory B cells, with tissue-specific higher proportions of unswitched memory cells in VAT and higher frequencies of switched memory B cells in the ovaries. These effects would suggest a regulated action by PCOS-derived IgG, altering energy metabolism which led to an immediate increase in body weight while concomitantly inducing a shift in both circulating and tissue-resident B cells towards altered cellular functionality. A metabolic shift could also further lead to a reprogramming of B cells towards a phenotype with a higher pro-inflammatory propensity as suggested by a recent study where peripheral B cells from women with PCOS were shown to have an increased capacity to produce TNF-α, which is attenuated by metformin treatment^26^. However, no variations were noted in the WT IgG-recipient mice metabolic phenotype, as glucose metabolism was not affected. This would be in line with the observations of Winer et al.^11^ already suggesting that preceding sterile inflammatory stimuli such as diet-induced pre-conditioning or induction of target autoantigens is required for proinflammatory IgG to have a critical role in rapid local and systemic metabolic changes. Such a stimulus affecting B cells frequencies may derive from direct or indirect testosterone/androgen receptor activation within the specific hyperandrogenic hormonal environment of PCOS.

Yet, although directly exposed to DHT before the onset of puberty, circulating B cell frequencies in the blood of peripubertal DHT-induced PCOS-like mouse model did not entirely reflect the distribution as seen in hyperandrogenic women. Additionally, we found a distinct profile of immunoglobulins in the peripheral blood of the peripubertal PCOS mouse model, with reduced levels of proinflammatory IgG2b and IgG2c titers. The increased concentration of circulating IgM in PCOS women was also not replicated. This raises the outstanding question regarding the influence of developmental origin and to what extent, if at all, does timing of androgen exposure alter B cell effector functionality in PCOS. Sensitivity of B cells to the influence of androgens is primarily during development as both pro- and pre-B cells within bone marrow express the AR, as do hematopoietic stem cells (HSC)^12^. Indeed, during fetal and early postnatal origin unique innate-like lymphocytes, such as Vg3+ dendritic epidermal T cells as well as so-called B-1a cells, and a fraction of innate-like marginal zone (MZ) B cells emerge which are not equally generated by adult bone marrow hematopoietic stem and progenitor cells (HSPCs)^27–30^. These cell types are characterized by distinct common features extending to both adaptive and innate immunity including their semi-invariant, often self-reactive antigen receptors, rapid responses to antigen stimulation and tissue resident localization^27^. Hence, the variations of observed IgM titers in PCOS women may be due to different B-1a cells frequencies, responsible for spontaneous testosterone-independent secretion of natural IgM antibodies (NAbs), which represent a large extent of circulating serum IgM^31^ that act with homeostatic housekeeping functions to multiple inflammatory reactions^32^. Although this remains purely speculative, as to whether this may be via immune activation related to the disease or a result of the disease driving the production of NAbs of IgM, we have previously shown that prenatal androgen exposure has severe effects on the health of offspring across generations, i.e. transgenerational transmission^17^, and alterations of IgM titers in lupus-prone (MRL/lpr) mice does induce a more severe autoimmunity^31^. Furthermore, while the focus of our findings was on the B cell lineage, early androgen exposure in ontogeny may have a similar impact on T cells and other innate lymphoid cells, with significant implications for the development of PCOS pathology. However, we also know that regulation of B cell number by testosterone/AR may occur also in mature B cells lacking AR independently of developmental effects as demonstrated in spleen from short-term castration of adult mice^33^. Within the context of sexual dimorphism in immunity and the effects of testosterone on B cells, previous research has focused on human male data or conditional AR knockout (ARKO) and castrated male mice. Both global ARKO models as well as castration cause a systemic alteration, which does not allow to distinguish in these studies if changes in the distribution of lymphocytes are due to a direct action of androgens on immature B cells expressing the AR or from secondary effects of surrounding tissue^34^.

A recent study has demonstrated how endogenous testosterone regulates indirectly mature splenic B cell number in a BAFF-receptor dependent manner via testosterone-mediated increase in sympathetic nervous transmission regulating BAFF-producing fibroblastic reticular cells (FRCs) in male mice^13^. There is however no data available regarding the concentration-response relation between BAFF levels, both local or systemic, and B cell numbers in mice. BAFF levels are overall tightly regulated and human studies suggest that serum BAFF levels within normal homeostatic ranges are inversely associated with peripheral B cell numbers^35,36^. This has led to the notion that estradiol accelerates autoimmunity while testosterone has an inhibiting action^37^. However, the singular setting of hyperandrogenic PCOS, with increased circulating and tissue-specific levels of testosterone along with higher estrogen-to-progesterone ratios leading to anovulatory cycles, as well as high estrogen levels during prenatal life, may disrupt the development of the thymus and its function in maintaining immune tolerance and are suspected to heighten autoimmune response in PCOS^38^. Indeed, serum levels of BAFF have been shown to be higher in women diagnosed with PCOS^14^. Our findings in the peripubertal DHT-induced PCOS mouse model showed metabolic alterations in glucose metabolism accompanied by variations among B cell populations, not only in blood-derived cells but also in immune cells seeding peripheral tissues. Trends of differential expression of complement receptor type 2 (CR2/CD21) on IgD^+^ naïve cells in both metabolic, immune, and reproductive tissues may suggest an effect deriving from BAFF overexpression affecting transitional B-cell maturation. BAFF is critical for the up regulation of expression of CD21 leading to greater proliferation and Ig secretion potential^39^, however, there is no direct link with the development of an autoimmune state. These effects were prevented and overall reduced by concomitant treatment with flutamide, supporting the idea that any modulation of B cells is regulated via AR activation. Finally, the lack of phenotype entirely with no variations neither in metabolic nor reproductive parameters in B cell reconstituted muMt^-^ mice as well as in IgG-recipient T cell-deficient RAG1 KO^-/-^ mice further suggest that the effects of B cells deriving from a PCOS-like environment may be dependent on androgen exposure of other leukocytes to fully promote impairment of metabolic parameters.

The fact that B cell depletion has profound effects on glucose homoeostasis is well accepted. For example, Rituximab, an anti-human CD20 mAb, used in the treatment of rheumatoid arthritis as well as B cell malignancies, can cause both hyperglycemia and severe hypoglycemia^40^. Depletion of B cells in mice with a CD20 mAb early in atherosclerotic disease has also shown therapeutic benefits in the abnormal glucose metabolism^41^. It is important to note that in these CD20 mAb therapies, the beneficial effects were linked to reduced T cell activation. In the present study, the lack of B cells did not prevent the development of a PCOS-like phenotype in muMt^-^ mice exposed to DHT, with both reproductive and metabolic effects, such as higher deposits of adipose tissue. Based on these results, in accordance with the lack of phenotype in RAG1 KO^-/-^ mice following IgG transfer, identification of the precise role for T cell activation in PCOS warrants further investigation as well as of tissue-resident immune cells such as macrophages, potentially skewing to a pro-inflammatory phenotype which may, in turn, be responsible for inflammatory cytokine production.

In conclusion, our study uncovers a previously unrecognized regulation via AR signaling, indirectly affecting B cell production of pathogenic IgG antibodies affecting energy metabolism. Moreover, our data raise a concern about uniquely identifying CD19^+^ B cells as a potential therapeutic target for PCOS as our findings do not support the notion that depletion of B cells is protective from developing PCOS-like traits. Concurrently increased levels of IgM may rather suggest a dual housekeeping function activity. Differences in the regulation of innate and adaptive immunity may be unique within the specific hormonal state of women with PCOS and should be investigated further.

## Material and Methods

### Human case–control explanatory study cohort

From September 2019 to March 2021, 15 women, all Caucasian ethnicity, were screened for a PCOS diagnosis at the Medical University Clinics in Graz (Austria) for either of the two main PCOS hyperandrogenic phenotypes: Phenotype A: Clinical hyperandrogenemia, oligoanovulation and polycystic ovarian morphology (PCOM) or Phenotype B: Clinical hyperandrogenemia and oligo-anovulation. Clinical hyperandrogenism was assessed using the modified Ferriman-Gallwey (FG) Score, with a self-reported score of eight or higher indicating hirsutism^42^. For total testosterone, a cut-off of 0.6 ng/mL (2.1 nmol/l) was used based on previously published data from a representative population sample^43^. Oligo-/anovulation was defined as menstrual cycles with a duration >35 days or the absence of menstruation for three or more consecutive months. Polycystic ovarian morphology, diagnosed by a gynecological ultrasound, was assessed based on medical history. Thyroid disorder, congenital adrenal hyperplasia, Cushing’s syndrome, hyperprolactinemia, androgen-secreting tumors, and pregnancy were excluded by laboratory measurements of thyroid-stimulating hormone (TSH), 17-hydroxyprogesterone (17OH-P), cortisol, prolactin, pregnancy test, and clinical examination. Exclusion criteria considered multiple factors affecting participants’ immune and hormonal profiles, such as neoplastic, infectious, and autoimmune diseases as well as currently used hormonal contraceptives or immunomodulating drugs. The final analyses of fasting blood samples were performed from a group of 15 women with PCOS and 22 controls were analyzed for B cell frequencies. Antibody variation was examined in the same women i.e. 15 PCOS and in 18 of the 22 controls. In the case of missing values due to, patients were excluded from the analysis report for that variable. All recruitment took place at the endocrinological Outpatient Clinic of the University Hospital Graz by routine doctors, and nurses involved in the project. All participants provided oral and written informed consent after a positive vote of the Ethics committee of the Medical University Graz (EK 31-560 ex 18/19). The work here described has been carried out in accordance with The Code of Ethics of the World Medical Association (Declaration of Helsinki) for experiments involving humans. For the extraction of IgG, from February 2020 to October 2020, 10 women accepted to voluntarily participate in the study. Seven women were diagnosed with PCOS, 1 did not fulfill the inclusion criteria and 2 decided to drop out, leaving 4 women with PCOS and 4 healthy donors were recruited. The diagnosis was conducted at the Medical University Clinics in Graz (Austria) according to the criteria.

### Clinical examination, blood sampling, and biochemical measurements

Anthropometric measures included weight, height, waist circumference, and body mass index (BMI) was calculated as weight (kg)/height (m)^2^ and waist-to-hip circumference. Baseline fasting blood samples were drawn for each participant in serum, EDTA, and lithium heparin tubes. Hormonal levels were assessed in fasting serum samples: total and free testosterone, androstenedione, and progesterone were measured by liquid chromatography–tandem mass spectrometry as described elsewhere^43^; sex hormone-binding globulin (SHBG), anti-Müllerian hormone (AMH), and insulin were measured by automated chemiluminescence immunoassay (ADVIA Centaur XP, Roche, Rotkreuz, Switzerland); serum luteinizing hormone (LH) and follicle-stimulating hormone (FSH) measured by enzyme-linked immunosorbent assay (ELISA, DIAsource Immunoassay, Belgium); Plasma total cholesterol, high-density lipoprotein (HDL) cholesterol, triglycerides, glucose were measured by automated enzymatic colorimetric assay (Cobas, Roche, Germany). The area under the curve (AUC) for glucose and insulin was calculated from the oGTT using the trapezoidal method. Serum SHBG and testosterone were used to calculate the free androgen index as serum testosterone/SHBG×100.

### Chemiluminescent ELISA

Chemiluminescent ELISA of human samples was performed as described elsewhere^44^ for total IgM, IgG and IgA. In brief, purified antihuman IgM, IgG and IgA (BD Pharmingen, San Jose, CA, USA) at concentrations of 5 μg/ml in 50 μl phosphate-buffered saline (PBS)-EDTA were added to each well of a 96-well white, round-bottom microtitration plate (MicrofluorII round-bottom; Thermo, Rochester, New York, USA) and incubated overnight at 4°C. After washing and blocking with Tris-buffered saline (TBS) with or without EDTA (pH 7.4, containing 1% bovine serum albumin (BSA), 30 min at room temperature), the plate was incubated with plasma samples in their respective dilutions in 1% BSA in TBS with EDTA (pH 7.4) for 2 h at room temperature or overnight at 4 °C. Alkaline phosphatase (AP)-labeled goat anti-human IgM (μ-chain specific; Sigma-Aldrich, Vienna, Austria; 1:50,000 in TBS BSA), AP-labeled goat anti-human IgG (γ-chain specific; Sigma-Aldrich, Vienna, Austria; 1:50,000 in TBS BSA) and AP-labeled goat anti-human IgA (α-chain specific; Sigma-Aldrich, Vienna, Austria; 1:50,000 in TBS BSA) was used for detection. AP-conjugated secondary reagents were detected using Lumi-Phos (Lumigen, Southfield, Michigan, USA; 33% solution in water) and a Synergy 2 Luminometer (BioTek, Winooski, Vermont, USA). Washing steps were performed on an ELx405 Select Deep Well Microplate Washer (BioTek, Winooski, Vermont, USA) with PBS or PBS-EDTA. Internal controls were included on each microtiter plate to detect potential variations between microtiter plates. The intra-assay coefficients of variation for all assays were 5–15%.

### Lymphocyte phenotyping of human samples

Blood samples from the baseline visit were processed within 4 hours for analysis by flow cytometry as previously described^45^. Briefly, for B-cell phenotyping, PBMCs were isolated from lithium heparin whole blood by Ficoll gradient density centrifugation. One million PBMCs were incubated with the following antibodies: CD19-VioGreen (clone REA675), anti-IgD-VioBlue (clone IgD26), CD27-APC (clone M-T271), CD86-PE-Vio770 (clone FM95), CD38-FITC (clone IB6), and anti-IgM-PE (clone PJ2-22H3, all purchased from Miltenyi Biotec, Bergisch Gladbach, Germany). Samples were measured using a FACSLyric flow cytometer (BD Biosciences, Franklin Lakes, NJ, USA). Data were analysed using the FACSSuite (BD Biosciences).

### Animals and study design

All mice experiments were carried out in compliance with the ARRIVE guidelines in accordance with the U.K. Animals (Scientific Procedures) Act, 1986 and associated guidelines, EU Directive 2010/63/EU for animal experiments. All animal experiments were approved by the Stockholm Ethical Committee for animal research (20485-2020) in accordance with the Swedish Board of Agriculture’s regulations and recommendations (SJVFS 2019:9) and controlled by Comparative Medicine Biomedicum at the Karolinska Institutet in Stockholm, Sweden. Mice were maintained under a 12-h light/dark cycle and in a temperature-controlled room with ad libitum access to water and a diet. All mice were on female on C57BL/6J background. For the transfer of human IgG 24 five-week-old female C57BL/6JRj mice were obtained from Janvier Labs. Rag1 KO^-/-^ were generated by breeding 10 Male and 10 Female B6.129S7-Rag1 (homozygous for Rag1) breeding pairs from Jackson Laboratory. For immune characterization of the peripubertal DHT-induced PCOS mice, 30 three-week-old female C57BL/6JRj mice were obtained from Janvier Labs ad left to acclimatize for 1 week. For the B cell reconstitution 10 three-week-old female C57BL/6JRj mice were obtained from Janvier Labs to develop the peripubertal DHT-induced PCOS model. This peripubertal DHT-induced PCOS mouse model was developed by implanting a 5 mm silastic implant containing 2,0-2,5 mg of continuously releasing DHT according to previously published protocol^20^, which was implanted subcutaneously in the neck region of 28-29 days old C57BL/6JRj female mice. Surgery was performed under light anesthesia with isoflurane. Control mice were implanted with an empty, blank implant. To investigate androgen receptor activation, a third group received, in addition to the DHT implant, a 4.5 mm continuously releasing pellet containing 25mg of flutamide (releasing time 90 days, Innovative Research of America, cat. number NA-152), an androgen receptor antagonist. Mice were randomly allocated to one of these three groups: control, DHT, DHT-flutamide. A PCOS-like phenotype was fully developed after 3 weeks of exposure. MuMt^-^ mutant mice were generated from 10 Male and 10 Female B6.129S2-Ighm (homozygous for Ighm) breeding pairs from Jackson Laboratory. No mice received further monthly implants.

### Purification and transfer of IgG

IgG from human sera was purified utilizing a HiTrap Protein G HP purification column (Bio-Sciences AB) according to vendors instructions. Briefly, samples were centrifuged at 3000 RCF for 5 minutes at 4°C and supernatant was diluted 5x with binding buffer. The final elution containing IgG was dialyzed overnight at 4°C against endotoxin-free PBS and further filtered to obtain sterile antibody solution. IgG concentration in each sample was measured by QUBIT (Thermo Scientific) according to vendor’s instructions and stored at −20°C. Samples from the serum of PCOS-affected women cohort or serum of the control group were separately pooled. The day before injection, samples were filtered and concentrated using Amicon Ultra-15 Centrifugal Filters, (30kDa MWCO - 15 mL sample volume) according to vendor’s instructions (Merck Millipore). Briefly, samples were thawed and kept at 4°C the night before concentration; after filtering samples through sterile 0.22 μm syringe filter, desired concentration was obtained by spinning multiple times at 1000g/4°C until reaching final concentration of 4 mg/ml of IgG antibody in a total volume of 450μl at injection day 1 and 3, and 365μl at injection day 10 of endotoxin-free PBS. Final IgG concentration was measured once again by QUBIT (Thermo Scientific). 7-week-old female C57BL/6JRj mice, randomly divided into 2 study-groups of 6 mice each, received purified human IgG (>98% pure) via intraperitoneal injection (i.p.) in endotoxin-free PBS on days 1, 3 and 10. To assess the role of T cells mediating the response to IgG, the same procedure was repeated utilizing age-matched in-house bred mutant Rag1 KO^-/-^ mice.

### Assessment of reproductive phenotype

In all groups, anogenital distance, a biomarker for androgen exposure, was measured at baseline and at sacrifice. For the transfer of human IgG, anogenital distance was measured 1 week after first i.p. injection in both WT and RAG1 KO^-/-^ mice. For immune characterization, anogenital distance was measured 3 weeks after DHT/flutamide implantation. For B cell reconstitution, anogenital distance in reconstituted muMt-mice was measured 2 weeks after reconstitution. Estrous cyclicity was assessed by daily vaginal smear for twelve consecutive days (three ovulatory cycles).

### Assessment of metabolic phenotype

Body weight development was recorded weekly. Body composition was assessed by magnetic resonance imaging (EchoMRI-100 system, Houston, TX, US) to measure total fat and lean mass in conscious mice. Glucose metabolism was measured by oGTT after a 5-h fast. Mice received 2 mg per gram body weight of D-glucose (20% glucose in 0.9% NaCl) administrated by orally by gavage. Blood glucose was measured at baseline and at 15-, 30-, 60- and 90-min following glucose administration (Free Style Precision). Blood was collected in EDTA coated capillary tubes at baseline and 15 min for insulin measurement by tail bleeding. Plasma separation is obtained by spinning the samples at 2000 g for 10 minutes at 4°C and stored at −20°C. Based on the study design for individual project objectives, for the transfer of human IgG as well as B cell reconstitution, mice were first assessed for oGTT when the expected effects from the transfer on glucose metabolism were most likely at their peak, followed Echo MRI evaluation. For project characterization of DHT-induced PCOS-like mouse model as well as the characterization of androgen exposed muMt-mouse model, mice were first screened through EchoMRI to measure total fat and lean mass and then subjected to an oGTT evaluation.

### Biochemical assessment of insulin and sex steroids in mice

Plasma insulin from oGTT was analyzed by an ELISA kit (Crystal Chem). Testosterone, androstenedione, estradiol, estrone, and progesterone were measured in serum using a high-sensitivity liquid chromatography–tandem mass spectrometry assay as previously described^46^.

### Tissue collection and cell isolation

For the transfer of human IgG, C57BL/6JRj WT mice were sacrificed at 13-14 weeks of age. Rag1 KO^-/-^ receiving human IgG were sacrificed at 16-17 weeks of age. For immune characterization of the peripubertal DHT-induced PCOS model, two independent experiments were conducted to evaluate separate timepoints: a first cohort of C57BL/6JRj mice were sacrificed at 20-22 weeks of age, while in a following assessment DHT-exposed C57BL/6JRj mice were sacrificed at 13-14 weeks of age. For the reconstitution of B^null^ muMt^-^ mice with splenic B cells following DHT exposure, a cohort of C57BL/6JRj mice were sacrificed at 8 weeks of age, 4 weeks after DHT implant, for the retrieval of spleen B cells. The B-cell reconstituted muMt^-^ mice were sacrificed at 11-12 weeks of age. For characterization of DHT-exposed MuMt^-^ model, mice were sacrificed at 13-14 weeks of age. All mice were sacrificed based on their ovulatory cycle stage in metestrous or diestrous, assessed by vaginal smears less than 2 hours prior sacrifice. Mice were fasted for 2 hours and anaesthetized with isoflurane (Isoflo vet, Orion Pharma Animal Health). Blood was drawn by cardiac puncture using a 21G needle; an aliquot of 150 μl was directly transferred to EDTA coated tube and placed on ice for FACS analysis. The remaining amount of blood was transferred to microvette capillary tubes (Sarstedt) for serum separation. After dissection, spleen and lymph nodes were kept on ice in phosphate-buffered saline without Ca^2+^ and Mg^2+^ (DPBS). Ovaries, endometrium, and VAT tissues were maintained in RPMI containing 2% FBS on ice for cell isolation. For analysis of sex steroid, serum in aliquots of 250 μl was separated by centrifugation at 5000 G for 10 min at 4°C.

### Comprehensive B lymphocyte phenotyping of mice tissues

To obtain single cells, spleen and inguinal and retroperitoneal lymph nodes were directly passed through a nylon wool sieve (100 μm cell strainer). After centrifugation at 300 RCF at 4°C for 5 min, erythrocytes (in spleen) were hemolyzed in 1 ml red blood cell lysis buffer (RBC lyse buffer; 0.16 M NH4Cl, 0.13 mM EDTA and 12 mM NaHCO3 in H2O), followed by a wash in 2 ml of FACS buffer (x2 the volume of RBC lysis). After a second centrifugation at 300 RCF at 4°C for 5 min, cells were resuspended in flow cytometry buffer (2% fetal bovine serum and 2 mM EDTA in PBS). Ovarian and uterus tissues were transferred into a 1ml and 3 ml digestive mix (1 mg/ml collagenase type I from 210U/mg, 0.8U DNase I, RPMI, 2% FBS) respectively, minced by fine scissors and digested by gentle shaking for 15 and 20 min respectively at 37°C. To inactivate the enzymatic activity, 2 ml and 6 ml respectively of cold flow cytometry buffer was added to ovaries and uterus and placed on ice before grinding tissues through a 100μm cell strainer. Samples were spun at 1000 G for 7 min at 4°C and resuspended in flow cytometry buffer. Visceral adipose tissue was minced by fine scissors in 5 ml digestive buffer based on RMPI containing 2% of FBS and 1 mg/ml collagenase type IV (type D, 0.15U/mg) and digested by gentle shaking for 20-25 min at 37°C. To inactivate the enzymatic activity, 10 ml of cold flow cytometry buffer was added and placed on ice before filtering suspensions through 100μm filter and further spinning at 500 G for 5 min. The resuspended cell pellet was left for 30 sec at RT in 500μl RBC lyse buffer, then washed in 1 ml of FACS buffer (x2 the volume of RBC lysis) and centrifuged at 500 G for 5 min at 4 °C. Blood volume of approximately 120 μl was placed twice in 1 ml of room temp RBC lysis buffer (for an approximate proportion of 1:10) for 5 and 2 minutes respectively, each time diluted in 2 ml of flow cytometry buffer and spun at 380 G for 5 min at 4°C. All tissue deriving cells were plated on 96-well round bottom plates and stained (Sarstedt, 83.3925.500) with FC-blocking antibody surface antigen staining (CD16/32, clone 2.4G2, BD Biosciences) diluted 1:100 in flow cytometry buffer, followed by incubation with the following antibodies: IgD-Pacific Blue (clone 11-26c.2a, BioLegend), CD19-BV480 or PE/Cyanine7 (clone 1D3, BD Biosciences or clone 6D5, BioLegend respectively), CD45R/B220-FITC (clone RA3-6B2, BD Biosciences), CD21/CD35-PE-CF594 (clone 7G6, BD Biosciences), CD138-PE/Cyanine7 (Syndecan-1, clone 281-2, BioLegend), CD27-APC (clone LG.3A10, BD Biosciences), IgM-APC/Cyanine7 (clone RMM-1, BioLegend), CD86-BV510 (clone GL1, BD Biosciences). Samples were measured using a FACS Canto II flow cytometer (BD Biosciences, Franklin Lakes, NJ, USA). Data were analyzed using FlowJo (BD Biosciences).

### Total antibody quantification in plasma by ELISA

Chemiluminescent ELISA was performed as described elsewhere^47^. Total IgM, IgG1, IgG2b, IgG2c, IgG3 and IgA antibodies in plasma were measured by ELISA. In brief, 96-well white round-bottomed MicroFluor microtiter plates (Thermo Lab systems) or immunoGrade, 96-well, PS Standard plates (781724; Brand) were coated with an anti-mouse IgM (Sigma; M8644; at 2 μg/ml), anti-mouse IgG1 (Biolegend; RMG1-1; at 2 μg/ml), antimouse IgG2b (BD Biosciences; R9-91; at 3 μg/ml), anti-mouse IgG2c (STAR135; at 1 μg/ml), anti-mouse IgG3 (BD Biosciences; R2-38; at 4 μg/ml) or anti-mouse IgA (BD Biosciences; C10-3; at 3 μg/ml) in PBS overnight and then washed three times with PBS and blocked with Tris-buffered saline containing 1% BSA (TBS/BSA) for 1 h at room temperature. Then wells were washed with either PBS (plates for IgM, IgG2b and IgG2c) or PBS supplemented with 0.05% Tween (plates for IgG1, IgG3 and IgA), and diluted mouse plasma was added in TBS/BSA to the wells and incubated overnight at 4 °C. Plates were washed, and bound Igs were detected with an antimouse IgM antibody conjugated to alkaline phosphatase (Sigma; A9688), the biotinylated forms of anti-mouse IgG1 (BD Biosciences; A85-1) or anti-mouse IgG2b (BD Biosciences; R12-3), anti-mouse IgG2c (JIR 115-065-208), anti-mouse IgG3 (BD Biosciences; R40-82) or anti-mouse IgA (BD Biosciences; C10-1). Wells were washed again as before and neutravidin conjugated to alkaline phosphatase was added where appropriate. Then, wells were washed again as before and rinsed once with distilled water, and 25 μl of a 30% LumiPhos Plus solution in dH2O (Lumigen Inc.) was added. After 75 min, the light emission was measured with a Synergy 2 luminometer (BIO-TEK) and expressed as RLU per 100 ms.

### Statistics

For statistical evaluation, Prism (version 9; GraphPad Software) and SPSS (version 28.0; SPSS) were used. All continuous data were screened for normality by Shapiro-Wilk test and equality of variance. Normally distributed data were compared using unpaired Student’s *t*-tests. Differences between more than two groups were determined by ANOVA followed by Tukey’s post hoc test. Differences were considered statistically significant at P<0.05. One patient or one animal was considered a biological replicate. In the case of missing values, patients or animals were excluded from the analysis for that variable. For the human monocentric casecontrol explanatory study, the sample size was calculated considering the distribution associated with specific PCOS phenotypes A and B. More than half of PCOS patients identified within the clinical setting demonstrate phenotype A, whereas the other three phenotypes (i.e., B, C, and D) have almost equal prevalence^48^, added to the observations that the presence of hyperandrogenism^49^, BMI^49^, and degree of menstrual irregularity^50^, while no ovarian morphology^51^, may be considered independent predictors of metabolic dysfunction. With the intention to evaluate the effects of double negative autoreactive B cells and assuming a predicted variation of 10% among total CD19^+^ B cell populations in PCOS patients based on previously reported assessments ^14^, the total sample of 40 subjects would achieve 90% power to detect differences among the means versus the alternative of equal means using an F test with a 0.05 significance level. The size of the variation in the means is represented by the effect size f = σm/σ, which is 0.31. Sample size was generated using PASS 15.0.6. For animal studies, no statistical methods were used to predetermine sample size. Animals were allocated to experimental groups arbitrarily without formal randomization. Investigators were not formally blinded to group allocation during the experiment.

## Data availability

Reserved DOI. At Mendeley Data: doi:10.17632/tcc2mbmys4.1.

## Acknowledgements

This work was supported by grants from Swedish Medical Research Council: project no. 2018-02435 and 2022-00550 (ESV); Novo Nordisk Foundation: Distinguished Investigator Grant – Endocrinology and Metabolism, NNF22OC0072904 (ESV) and NNF19OC0056647 (ESV); Diabetes Foundation: DIA2021-633 and DIA2022-708 (ESV); Strategic Research Program in Diabetes at the Karolinska Institutet (ESV); Karolinska Institutet KID funding: 2020-00990 (ESV); Regional Agreement on Medical Training and Clinical Research between the Stockholm County Council and the Karolinska Institutet: 20190079 (ESV); EMBO Scientific Exchange Grants 2021: STF 8938 (AA); European Research Council (ERC) under the European Union’s Horizon 2020 research and innovation program under the grant agreement no. 866075 (CIS); Knut and Alice Wallenberg Foundation no. 018.0161 (CIS); Austrian Science Fund (FWF) project number W1241 (BOP).

## Author contributions

Conceptualization: AA; ESV; BOP; MHS. Methodology: AA; ST; ESV. Investigation: AA; ST; SR; HL; GE; CL; ST; JM; KS; CO. Data acquisition, analysis, and visualization: AA; ST; MHS; ESV. Project administration: AA; ESV. Supervision: ESV; BOP; CIS; MK. Wrote the manuscript: AA; ST; ESV. All authors: reviewing and editing the manuscript. Funding acquisition: AA; ESV; BOP; CIS

## Supplementary

**Fig. S1.**
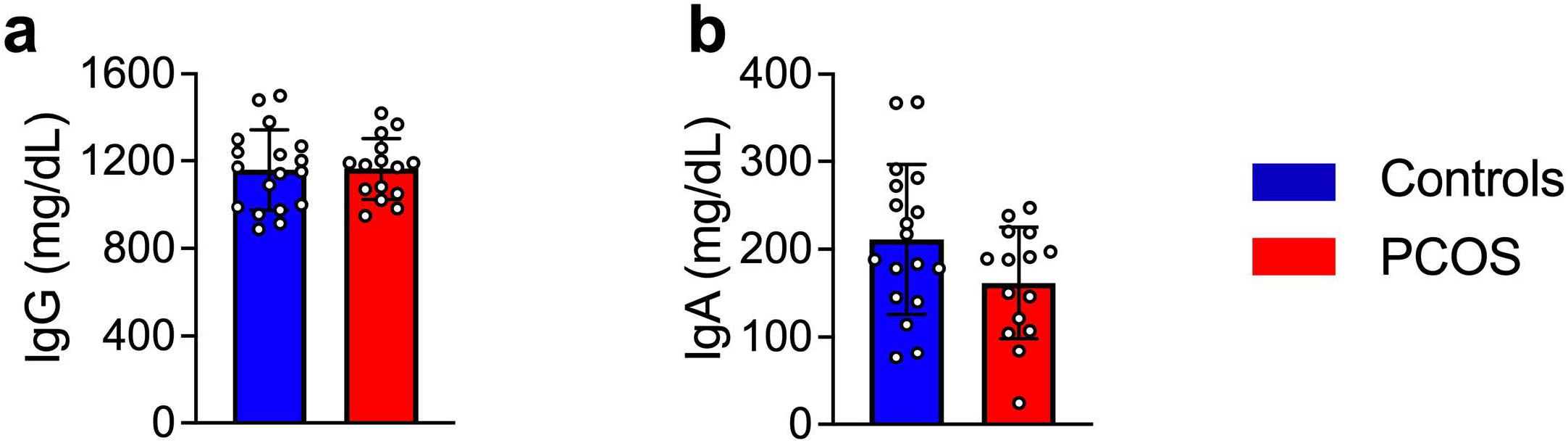
Circulating antibody titres in women with and without PCOS. **a.** Circulating IgG titres **b.** Circulating IgA titres. Controls n=18 PCOS n=15. All bars indicate means, circles represent human individuals. Unpaired Student’s t-test for analysis of antibody titres. *P<0.05, **P<0.01, ***P<0.001.

**Fig. S2.**
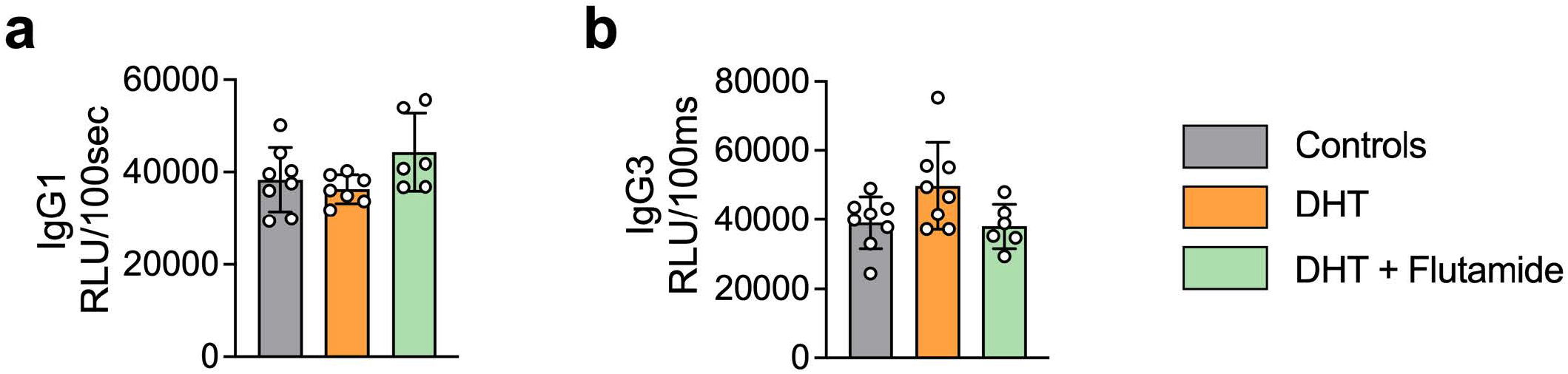
Circulating antibody titres in DHT-induced PCOS-like mouse model. **a.** Circulating IgG1 titres Controls n=8 DHT n=7 Flutamide n=6 **b.** Circulating IgG3 titres. Controls n=8 DHT n=8 Flutamide n=6. All bars indicate means, circles represent individual mice. Unpaired Student’s t-test for analysis of antibody titres. *P<0.05, **P<0.01, ***P<0.001.

**Supplementary Table 1 (S1).** Clinical characteristics of women with polycystic ovary syndrome (PCOS) and women without the syndrome used for characterization of main B cell lineages and subpopulations based on pan B cell surface marker CD19.

|  | **Controls**<br>**(n=22)** | **PCOS**<br>**(n=15)** | **P value** |
| --- | --- | --- | --- |
| Age (years) | 36.3 (21-50) | 26.4 (24-38) | **0.003** |
| *Anthropometry* |  |  |  |
| BMI (kg/m<sup>2</sup>) | 22.18 (18.31-32.15) | 24.39 (19.16-39.89) | 0.078 |
| Waist-to-Hip-Ratio | 0.8 (0.74-0.91) | 0.82 (0.71-0.92) | 0.387 |
| *Endocrine measure* |  |  |  |
| Free Testosterone (ng/mL) | 0.25 (0.06-0.49) | 0.43 (0.02-0.88) | **0.015** |
| Total Testosterone (ng/mL) | 0.86 (0.8-1.74) | 1.49 (0.07-3.05) | **0.019** |
| Free Androgen Index (FAI) | 1.2 (0.22-2.48) | 2.6 (0.11-20.34) | **0.021** |
| Androstenedione (ng/mL) | 2.8 (1.17-5.99) | 3.9 (2.27-6.53) | **0.002** |
| *Metabolic measures* |  |  |  |
| Cholesterol (mg/dL) | 177 (128-217) | 180 (135-214) | 0.597 |
| HDL (mg/dL) | 67 (46-87) | 51 (36-90) | **0.001** |
| LDL (mg/dL) | 97.3 (45.8-115.8) | 108.6 (78.6-160.4) | **0.049** |
| Triglycerides (mg/dL) | 65.5 (44-97) | 82 (56-124) | **0.008** |
| Glucose (mg/dL) | 88 (76-104) | 91 (77-111) | 0.452 |

Data are median ± range. Comparisons between groups were made using T-test. HDL. high density lipoprotein; LDL. low density lipoprotein.

**Table S2.** Clinical characteristics of IgG donors, women with polycystic ovary syndrome (PCOS) and women without the syndrome.

|  | **Controls**<br>**(n=4)** | **PCOS**<br>**(n=4)** | **P value** |
| --- | --- | --- | --- |
| Age (years) | 27 (22-31) | 25 (23-35) | 0.943 |
| *Anthropometry* |  |  |  |
| BMI (kg/m<sup>2</sup>) | 26 (19.4-29.8) | 25 (21.3-28.2) | 0.909 |
| Waist-to-Hip-Ratio | 1 (0.79-0.91) | 1 (0.70-0.91) | 0.515 |
| *Endocrine measure* |  |  |  |
| LH (mU/mL) | 7 (4.40 – 11.40) | 10 (5.19-38.60) | 0.351 |
| FSH (mU/mL) | 4 (2.74 – 6.91) | 7 (5.59 – 8.61) | **0.058** |
| Progesterone (ng/mL) | 10 (0.20 – 13.60) | 1 (0.60 – 1.05) | 0.075 |
| Free Testosterone (ng/mL) | 2 (0.84 – 2.68) | 3 (0.29 – 3.03) | 0.939 |
| Total Testosterone (ng/mL) | 0 (0.17 – 0.40) | 0 (0.30 – 0.40) | 0.107 |
| Androstendione (ng/mL) | 3.11 (1.21 – 4.56) | 4 (1.98 – 4.69) | 0.514 |
| SHBG (nmol/L) | 63 (52.8 – 88.8) | 62 (43.9 – 105) | 0.952 |
| Free Androgen Index (FAI) | 0 (0.3 -0.6) | 1 (0.5 – 0.7) | 0.107 |
| AMH (ng/mL) | 4 (2.40 – 4.66) | 8 (4.97 – 9.96) | **0.011** |
| *Metabolic measures* |  |  |  |
| Cholesterol (mg/dL) | 160 (132 – 184) | 153 (146 – 172) | 0.801 |
| HDL (mg/dL) | 77 (42 – 80) | 62 (49 – 71) | 0.458 |
| LDL (mg/dL) | 75 (66 – 89) | 74 (67 – 107) | 0.691 |
| Tryglicerides (mg/dL) | 71 (43 – 91) | 74 (68 – 82) | 0.632 |

Data are median ± range. Comparisons between groups were made using T-test. HDL. high density lipoprotein; LDL. low density lipoprotein.

## Appendix

**Table A1.** Antibodies for total IgM, IgG and IgA titring and lymphocyte phenotyping of human samples.

| Commercial reagent | Clone | Supplier | Catalog # | RRID |
| --- | --- | --- | --- | --- |
| anti-human IgM | G20-127 | BD Pharmingen | 555780 | AB_396115 |
| anti-human IgG | G18-145 | BD Pharmingen | 555784 | AB_396119 |
| anti-human IgA | G18-1 | BD Pharmingen | 555886 | AB_396198 |
| Goat Anti-Human IgM, μ-Chain Specific Alkaline Phosphatase Conjugate | | Sigma-Aldrich, Vienna, Austria | 401902 | N/A |
| Goat Anti-Human IgG Antibody, Alkaline Phosphatase conjugate |  | Sigma-Aldrich, Vienna, Austria | AP112A | N/A |
| Goat Anti-Human IgA (α-chain specific)-Alkaline Phosphatase | | Sigma-Aldrich, Vienna, Austria | A9669 | N/A |
| CD19 Antibody, anti-human, VioGreen | REA675 | Miltenyi Biotec | 130-113-649 | AB_2726202 |
| Anti-IgD-VioBlue antibody | IgD26 | Miltenyi Biotec | 130-099-484 | AB_2659773 |
| CD27-APC, human antibody | M-T271 | Miltenyi Biotec | 130-093-186 | AB_1036201 |
| CD86-PE-Vio770, human antibody | FM95 | Miltenyi Biotec | 130-113-573 | AB_275113 |
| CD38-FITC, human antibody | IB6 | Miltenyi Biotec | 130-092-259 | AB_615091 |
| Anti-IgM-PE antibody | PJ2-22H3 | Miltenyi Biotec | 130-093-075 | AB_1036088 |

**Table A2.** Antibodies for lymphocyte phenotyping of mouse samples.

| Commercial reagent | Clone | Supplier | Catalog # | RRID |
| --- | --- | --- | --- | --- |
| BV480 Rat Anti-Mouse CD19 | 1D3 | BD Biosciences | 566107 | AB_566107 |
| PE/Cyanine7 anti-mouse CD138 (Syndecan-1) Antibody | 281-2 | BioLegend | 142514 | AB_2562198 |
| Pacific Blue™ anti-mouse IgD Antibody | 11-26c.2a | BioLegend | 405712 | AB_405712 |
| FITC Rat Anti-Mouse CD45R/B220 | RA3-6B2 | BD Biosciences | 553087 | AB_394617 |
| PE-CF594 Rat Anti-Mouse CD21/CD35 | 7G6 | BD Biosciences | 563959 | AB_2738511 |
| APC Hamster Anti-Mouse CD27 | LG.3A10 | BD Biosciences | 560691 | AB_1727455 |
| APC/Cyanine7 anti-mouse IgM Antibody | RMM-1 | BioLegend | 406516 | AB_10660305 |
| BV510 Rat Anti-Mouse CD86 | GL1 | BD Biosciences | 563077 | AB_2737991 |
| PE/Cyanine7 anti-mouse CD19 Antibody | 6D5 | BioLegend | 115520 | AB_313655 |

**Table A3.**
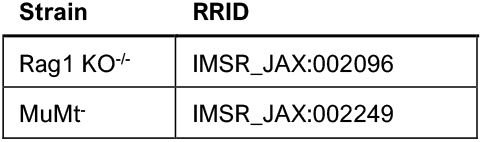
Transgenic mouse models.

